# Allosteric Role of Substrate Occupancy Toward the Alignment of P-glycoprotein Nucleotide Binding Domains

**DOI:** 10.1101/358549

**Authors:** Lurong Pan, Stephen G Aller

## Abstract

P-glycoprotein (Pgp) is an ATP-binding cassette transporter that eliminates toxins from the cell but causes multidrug resistance in chemotherapies. The crystal structures of Pgp revealed drug-like compounds bound to an inward-facing conformation in which the energy-harnessing nucleotide binding domains (NBDs) were widely separated with no interfacial interaction. Following drug binding, inward-facing Pgp must transition to an NBD dimer conformation to achieve ATP binding and hydrolysis at canonical sites defined by both halves of the interface. However, given the high degree of flexibility shown for this transporter, it is difficult to envision how NBDs overcome entropic considerations for achieving proper alignment in order to form the canonical ATP binding site. We explored the hypothesis that substrate occupancy of the polyspecific drug-binding cavity plays a role in the proper alignment of NBDs using computational approaches. We conducted twelve atomistic molecular dynamics (MD) simulations (100-300 ns) on inward-facing Pgp in a lipid bilayer with and without small molecule substrates to ascertain effects of drug occupancy on NBD dimerization. Both apo- and drug-occupied simulations showed NBDs approaching each other compared to the crystal structures. Apo-Pgp reached a pseudo-dimerization in which NBD signature motifs for ATP binding exhibited a significant misalignment during closure. In contrast, occupancy of three established substrates positioned by molecular docking achieved NBD alignment that was much more compatible with a canonical NBD dimerization trajectory. Additionally, aromatic amino acids, known to confer the polyspecific drug-binding characteristic of the internal pocket, may also govern polyspecific drug access to the cavity. The enrichment of aromatics comprising the TM4-TM6 portal suggested a preferential pathway over the aromatic-poor TM10-TM12 for lateral drug entry from the lipid bilayer. Our study also suggested that drug polyspecificity is enhanced due to a synergism between multiple drug-domain interactions involving 36 residues identified in TM1, 5, 6, 7, 11 and 12.

**Author Summary:** P-glycoprotein (Pgp) is an active drug pump known to cause clinical multi-drug resistance. The static atomic structure of Pgp was determined by trapping an inward-facing conformation bound to small molecule substrates by crystallization, however the effect of substrates on Pgp dynamics following binding is poorly understood. In this study, six apo-Pgp and six drug-occupied Pgp were simulated using unconstrained atomistic molecular dynamics (MD) for 100-300 ns. We demonstrate an allosteric communication of drug binding “from the top down”, that is from the TMDs to the NBDs that promotes NBD alignment and trajectories that favor canonical ATP binding. Other analyses suggested that aromatic amino acids in both the central drug-binding cavity and the “front portal” (TM4/TM6) confer polyspecific recognition. Additionally, comparison of the thermal B-factors between the experimental measurement and MD simulation indicated that different physical and chemical environments (temperature, *in surfo* vs. *in meso*, solution compositions) only alter the regional scales of thermal fluctuations but not the patterns of these motions. Lastly, DCCM and normal mode analyses were used to decipher thermal motions and the motion correlations between various domains in Pgp, allowing us to propose a substrate allosteric mechanism and an energy conservation mechanism during the catalytic cycle.

## Introduction

P-glycoprotein (Pgp) ^1^ is a transmembrane drug-efflux pump that belongs to the ATP-binding cassette (ABC) transporter superfamily ^2^. Pgp eliminates small molecules including anti-cancer drugs, herbicides, fungicides and other xenotoxins 3-5 by capturing them within the lipid bilayer and pumping them out of cells ^4^. The hallmark of Pgp function is the recognition and transport of a large number of diverse molecules making it one of the most polyspecific transporters in the superfamily ^6-12^. Experiments using endothelial cells of the blood-brain barrier in the knockout mouse, *mdr1a(-/-)*, demonstrate a critical role of Pgp in protecting the brain from toxins ^13^. Pgp serves as a protection mechanism against harmful chemicals, but overexpression can result in multidrug resistance to cancer chemotherapies.

The x-ray crystal structures of Pgp ^14,15^ provided the first snapshots at atomic resolution for understanding its polyspecificity and catalytic mechanisms. The structure of Pgp has two transmembrane domains (TMDs) with twelve helices (TM1-12) and two nucleotide-binding domains (NBDs), which comprise canonical ABC signature motifs. Pgp was crystallized in “inward-facing” conformations in the absence of presence of drug-like cyclic peptides which revealed the large internal polyspecific drug binding cavity. The volume of the internal hydrophobic cavity of Pgp within the lipid bilayer is currently the largest described for any protein (~ 6,000 Å^3^) and was shown biochemically to be capable of binding at least two transported substrates simultaneously ^14,16^. The composition of residues in the drug binding cavity is mostly hydrophobic and comprises a group of residues, some of which are represented by strong electron density in the mouse Pgp crystal structure, and some lack density indicating disordered side chain positioning. The complete transport cycle is known to include large conformational changes that couple the drug binding sites in the TMDs ^17,18^ to the dimerization of the NBDs upon binding nucleotide ^19,20^. Pgp utilizes the energy from ATP binding and hydrolysis in the NBDs to catalyze the translocation of drugs across the TMDs in the lipid bilayer.

Thermodynamic and kinetic effects of drugs, ATP, lipids and water on the protein cannot be ascertained from inspecting crystal structures alone. Conventional MD simulations ^21-24^ on Pgp in the inward-facing conformation have been conducted previously as well as targeted steered MD simulations between inward-facing and outward-facing conformations ^25,26^. These studies provided valuable insight regarding Pgp catalytic mechanisms. However, the previous simulations were based on the original crystal structures that have since been shown to contain registry errors and other problems ^15^ which may have produced artifacts. The simulations might involve various artifacts as a result of the problematic starting structures^21-24,26^. Previous MD simulations also characterized ATP binding of the inward-facing conformation ^21-23^ even though this conformation is generally considered to have low affinity for ATP. Allosteric communication between the NBDs and TMDs has not been extensively explored in previous MD studies. Docking experiments were performed with various drug substrates in the TMD substrate-binding cavity ^22,27^ but MD was not used to examine the dynamic effects these substrates might have on conformational change. Steered MD was performed in the 10 ns ^26^ and 50 ns _25_ timescales, which are much shorter compared to the μs+ timescale expected for sufficient sampling of the rather large conformational changes expected for alternating access transport processes. Furthermore, targeted MD simulations that apply symmetric forces on Pgp may overlook important biological insights due to the asymmetric nature of the transporter. Specifically, the amino acid sequences of the two pseudosymmetric “halves” of mouse Pgp only share 59.4% similarity and experiments 28-30 have established that ATP binding/hydrolysis alternates, implying very asymmetric conformational changes of the protein occur during catalytic cycling. Computational studies of Pgp function must therefore take the asymmetric architecture of Pgp and what is already known about Pgp biochemistry into account.

To date, the sequence-structure-function relationships of Pgp have not been fully understood, but when characterized in more detail, would likely provide a broad framework for understanding polyspecific drug transport dynamics of many transporters in the ABC superfamily. Furthermore, the allosteric communication of drug binding from the TMDs to the NBDs of Pgp has not been adequately characterized. Nor has the energetic and functional consequences of the pseudo-symmetric two halves of mammalian Pgp, as opposed to symmetric dimers found in earlier metazoans or protozoans, been explored.

Our present work employed computational approaches including drug docking and MD simulations to address gaps in understanding allosteric mechanisms in Pgp. We used the substantially improved inward-facing Pgp crystal structures _15_ as starting structures for the MD simulations. The structures are stable by unconstrained all-atom MD simulations in contrast to the original structures that were previously shown to be unstable by MD ^31^. Six apo-Pgp and six drug-occupied Pgp structures were simulated using unconstrained atomistic MD for 100 ns to 300 ns with a system size of 209K atoms which allowed sufficient sampling of microstates and longer timescales compared to previous MD simulations on inward-facing Pgp. The study demonstrated detailed allosteric regulation of drug binding in the TMDs on the NBDs, and an accompanying asymmetric structural dynamics. Differential effects between lipid and drug binding on the conformational change of TMDs and NBDs were observed. NBD alignment that favors a canonical ATP binding conformation is promoted allosterically by drug occupancy, but drug-free Pgp (apo-Pgp) fails to achieve proper alignment in the timeframe of the simulations resulting in a pseudo-dimerization from a counter-clockwise twisting of NBDs also observed in simulations ^21,31^. The drug entry pathway and the poly-specific drug-binding network were also examined. The structural bases of several distinctive features of mouse Pgp sheds light on: 1) the drug translocation pathway from lipid bilayer to internal cavity, 2) the asymmetric allosteric regulation between TMDs and NBDs by two halves and 3) polyspecificity of the drug binding cavity. Normal mode analysis illustrated various thermal motions promoted or inhibited by drug binding. In addition, crystallization in detergents, i.e. *in surfo* crystallization, appears to result in protein conformations that are much less frequently sampled compared to conformations sampled in a lipid bilayer, i.e. *in meso*. Based on our findings, we propose a substrate-mediated allosteric mechanism and an energy conservation mechanism in the absence of drugs.

## Results and Discussion

### Rationale for Simulation Redundancy Setup

A total of twelve models with six apo-Pgp and six drug-Pgp models were thus simulated in this study as summarized in (Table 1). The protein was properly placed in the lipid bilayer by matching hydrophobic regions between protein and lipid and using tryptophan residues on the protein as anchor (Fig S1).

**Table 1.** Starting Models Used for MD simulations

| Models | Abbreviation | Crystal structure | Drug | MD |
| --- | --- | --- | --- | --- |
| 1 | 4M1M (A, apo) | 4M1M molecule A | none | 300 ns |
| 2 | 4M2S (A, apo) | 4M2S molecule A | removed | 100 ns |
| 3 | 4M2T (A, apo) | 4M2T molecule A | removed | 100 ns |
| 4 | 4M1M (B, apo) | 4M1M molecule B | none | 100 ns |
| 5 | 4M2S (B, apo) | 4M2S molecule B | removed | 100 ns |
| 6 | 4M2T (B, apo) | 4M2T molecule B | removed | 100 ns |
| 7 | 4M2S (A, QZ59-RRR) | 4M2S molecule A | cyclic-tris-(R)-<br>valineselenazole<br>(QZ59-RRR) | 300 ns |
| 8 | 4M2S (B, QZ59-RRR) | 4M1M molecule B | cyclic-tris-(R)-<br>valineselenazole<br>(QZ59-RRR); | 100 ns |
| 9 | 4M2S (A, QZ59-RRR(S)) | 4M2S molecule A | cyclic-tris-(R)-<br>valinethiazole<br>(QZ59-RRR(S)) | 100 ns |
| 10 | 4M2S (B, Irinotecan) | 4M2S molecule B | Irinotecan (docked) | 100 ns |
| 11 | 4M2S (A, Vinblastine) | 4M2T molecule A | Vinblastine (docked) | 100 ns |
| 12 | 4M2S (A, Etoposide) | 4M2T molecule A | Etoposide (docked) | 100 ns |

Three corrected Pgp crystal structures (PDB codes 4M1M, 4M2S and 4M2T) yielded six slightly different initial conformations, with the greatest differences embodied by the two molecules of the asymmetric unit called molecule A (mol A) and molecule B (mol B). The root-mean-square deviations (RMSD) between mol A and mol B were 3.34 Å for 4M1M, 3.19 Å for 4M2S and 3.06 Å for 4M2T. Using mol A and mol B of 4M1M as the reference set, the RMSDs were much smaller when compared to the counterpart conformations of the other two crystal structures (0.549 Å for 4M2S-mol A, 0.452 Å for 4M2T-mol A, 0.683 Å for 4M2S-mol B and 0.659 Å for 4M2T-mol B). The exercise highlights the fact that mol A and mol B are distinct conformations of Pgp that do not significantly vary between apo- and drug-bound forms. Therefore, the simulations of all six structures are essentially equivalent to triplicate simulations of each of two distinct conformations of the asymmetric unit. The drug-occupied model 4M2S (mol A and mol B) has two protein conformations with the same drug QZ59-RRR located in two slightly different but overlapping regions of the binding cavity. The use of two slightly different starting structures increases the sampling size and reduces the risk of bias associated with using a single initial conformation. Additional drugs were simulated on both mol A and mol B to test the effects of different drugs on the equilibrated conformations. The average results of simulating the six different drug-occupied Pgp structures provided insights into the mechanism of polyspecificity.

All simulations essentially reached equilibrium as measured by a plateau of C_α_ RMSD achieved at ~50 ns (Fig. S2), and at equilibrium, apo-Pgp exhibited higher conformational fluctuation than that of any drug-occupied Pgp. Initially, the drastic change in RMSDs compared to the initial structure were primary cause by the dimerization process of NBDs and flexible nature of NBDs at the inward facing conformations. However, only minor oscillations in RMSD were observed after 50 ns, which indicated that the protein achieved a set of meta-stable states of similar energies with relatively stable inter-domain configuration in equilibrium. The equilibrated trajectories (2000 frames) of the last 20 ns (80 ns– 100 ns) were used to obtain ensemble averages at equilibrium. 300 ns simulations were performed for two species, 4M1M (A, apo) and 4M2S (A, QZ59-RRR), to assure that the conclusions based on the shorter simulations were not affected by the time scale of the simulations. The RMSD (Fig. S2) and other analyses showed that the equilibrated confirmations of both species were achieved by the end of 100 ns and no significant differences were apparent between the trajectories in the final 20 ns of the 100 ns simulations and the last 200 ns of the longer simulations.

### B-factors between *in surfo* crystallization and *in meso* MD simulation

A potential concern of membrane protein crystallization in detergent without a lipid bilayer (*in surfo*) is that the resulting structures may yield different properties (protein conformations, atomic displacements, etc) compared to properties resulting from the physiological environment afforded by the lipid bilayer (*in meso*). In order to directly compare the *in meso* simulations with *in surfo* structures, root-mean-square fluctuations (RMSF) of the equilibrated structures were calculated and converted to B-factors to compare with the experimentally derived B-factors of the mouse Pgp crystal lattice 15 (Fig. 1, Fig. S3). The B-factors for the simulations were calculated using the same function (B=8π^2^RMSF^2^) as is used by the algorithm in Phenix ^32^ that was used to refine the corrected mouse Pgp crystal structures 15. TM helices in direct contact with lipids (Fig. 1) exhibited smaller B-factors in the simulations compared to that of the crystal lattice, likely a property conferred by the bilayer. However, the regions exposed to solvent in the simulations including NBDs and extracellular loops (ECLs) and intracellular helices (IHs) had larger B-factors compared to the same regions located in the crystal structure most likely due to the higher temperature (310 K simulations vs. 277 K crystallization), and/or less constraint from a lack of crystal packing in the simulations. It is noteworthy that the B-factors were affected by crystal lattice packing such that NBD1, TM2, 11 showed slightly smaller regional B-factors compared to their pseudo-symmetrical counterparts due to inter-molecular packing rather than a symmetric packing by the lipid bilayer (Fig. S3).

**Fig 1.**
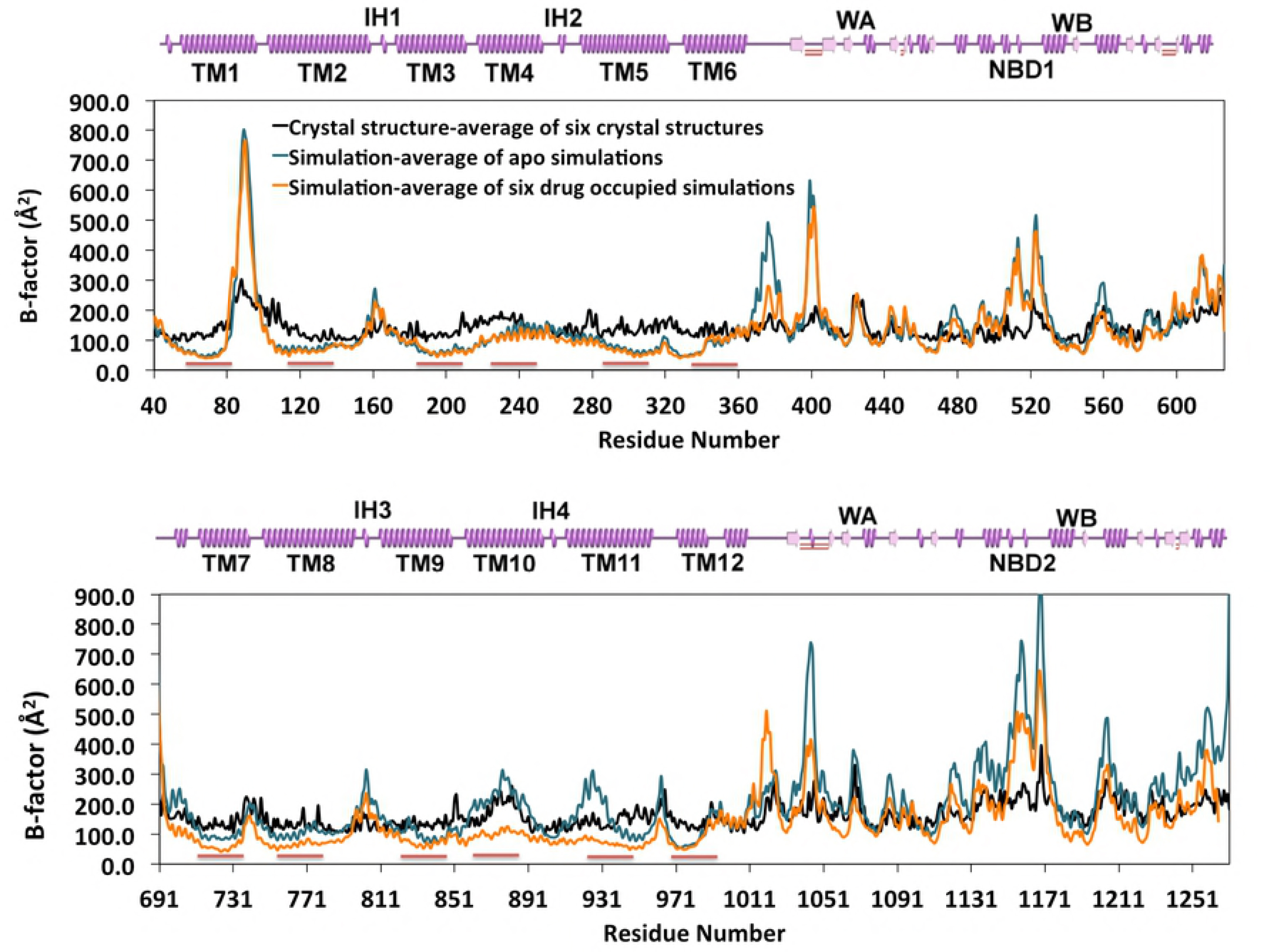
B-factors (Å^2^) of protein C_α_. The B-factors of crystal structures are from PDB 4M1M. The B-factors (B=8π^2^RMSF^2^) of simulations are measured based on the RMSFs of C_α_ from trajectories of the last 20 ns simulations and are averaged over six simulations for apo-Pgp (as blue lines) and drug-occupied Pgp (as orange lines). Red bars under each graph represent the lipid binding regions of Pgp. 2D cartoon representatives of the secondary structures are in pink. Domains are shown in the same sequence order as that of the RMSF (helix as spring, coil and turn as straight line, beta sheet as arrow). The Figs of the two pseudo-symmetric halves (half1: residue 40-626, half2: residue 691-1271) are displayed in parallel for comparison.

The analysis of B-factors also shows the asymmetry between two halves of Pgp. In apo-Pgp, TM10, 11 and NBD2 has larger fluctuation than that of their pseudo-symmetric counterparts TM4, 5 and NBD1. In addition, ECL1 between TM1 and TM2 has larger fluctuation than that of ECL3 between TM7 and TM8. Drug occupancy (Fig. 1, orange curve) did not significantly alter the fluctuation of “half1” but altered “half2” in the TMDs and NBDs. Since TM10 and TM11 are mainly lipid binding domains as oppose to drug binding domains based on crystal structure, the results indicate that drug binding alters the manner in which lipids interact with these TMs. The B-factors of Walker B of NBD2 was also reduced by drug binding, which indicates that drug binding also affects the crosstalk between NBD2 and IH2.

Interestingly, the peaks and valleys of the three sets of B-factor data (Fig. 1, black, blue and orange curve) exhibit a regular matching oscillation pattern. The patterns are also consistent in simulations of outward-facing Pgp ^33^. These data indicate that different physical and chemical environments (temperature, *in surfo* vs. *in meso*, solution compositions) only alter the regional scales (amplitudes) of thermal noise but not thermal patterns (frequencies). The results suggest that B-factors are intrinsic to the protein and may not be significantly influenced on a global level from the experimental conditions. Measurement of B-factors in MD simulations thus provides a practical way to examine the integrity of a crystal structure and the approach used to refine the structure.

### Secondary structure stability and conformational change

The secondary structure stability was measured over time (Fig. S4). All 12 runs demonstrated a high degree of secondary structure stability (consistent folded helix and beta sheet components) across the entire protein for the entire duration of the simulations and are consistent with the refined crystal structure15. The results were also supported by biochemical studies that showed that conformational change of Pgp during drug transport involves only tertiary structural changes whereas secondary structure remains essentially unchanged ^30,34,35^.

The conformational change was monitored using landmark residue pair distances between two pseudo-symmetric halves (Fig. 2). The results revealed a very different conformational change trajectory between apo-Pgp and drug-occupied Pgp. The results were also compared with the outward-facing Pgp with both MgATP^2−^ bound^33^. Due to the timescale limitation of conventional MD simulations, the conformational changes observed in these simulations (100~300 ns) could not achieve the complete transformation (~μs) from inward facing to the outward facing conformation. But in general, the drug-occupied Pgp simulations resulted in a more similar conformation towards the outward-facing structure compared to that of the apo-Pgp simulations. Specifically, the difference between the apo and drug-occupied simulations was insignificant at TMDs. The simulations showed a slight opening of “extracellular gates” (A79-T736, G325-E968 and G207-G850) compared to the inward-facing crystal structure but not as wide open as the outward-facing structure. Considerable closure of the drug-binding cavity was observed in both apo and drug-occupied simulations, especially for a landmark pair near the bottom of the inner leaflet (Q343-Q986). The surrounding lipid bilayer in the simulations might be the cause of such closure compared to that of crystal structure.

**Fig 2.**
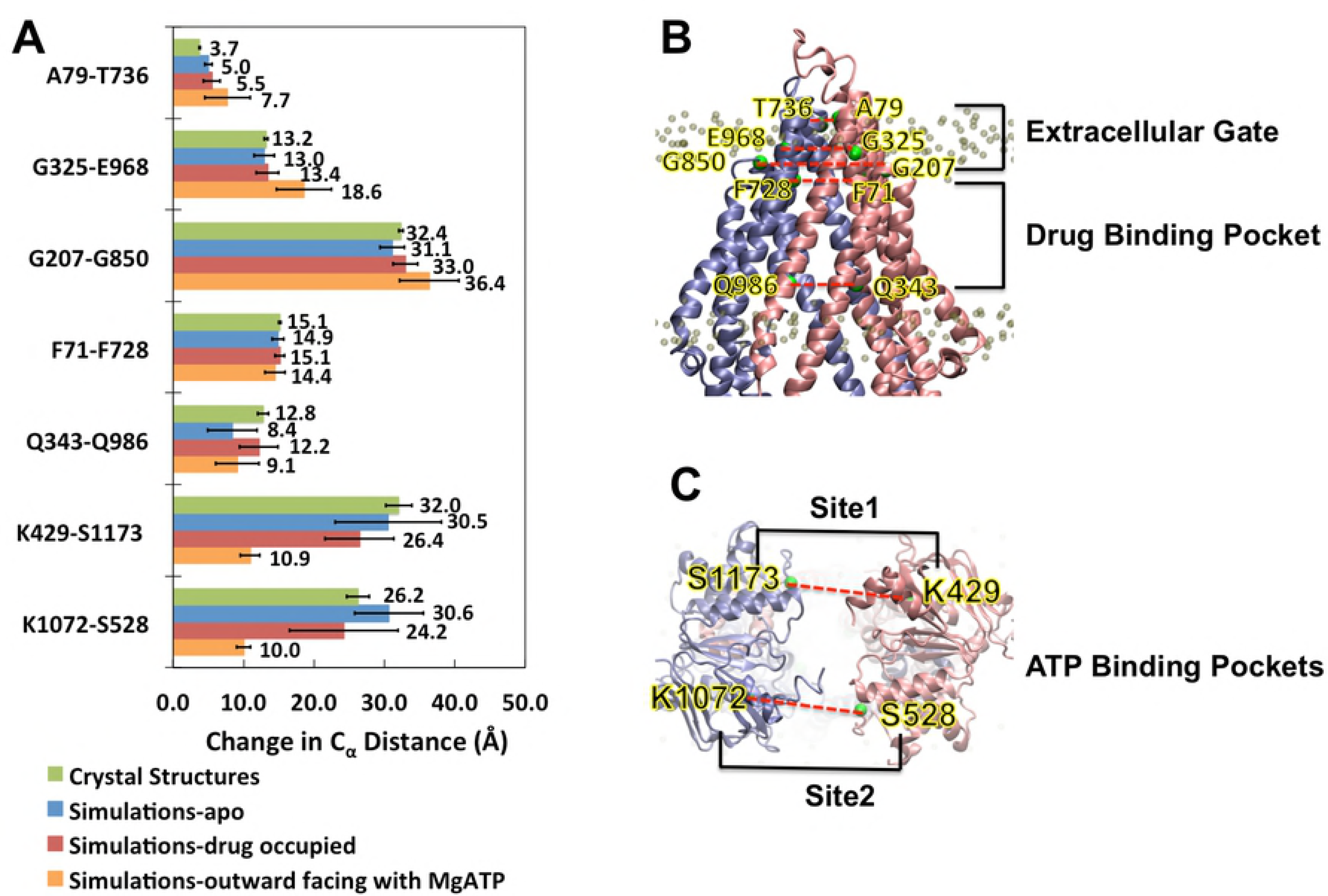
Conformational change monitored by landmark residues. In panel A, change in pairwise distances of C_α_ during simulations is measured from trajectories of the last 20 ns. The average value of 6 crystal structures (PDB: 4M1M(mol A, mol B), 4M2S (mol A, mol B) and 4M2T (mol A, mol B)). The average of three 200 ns Pgp-MgATP simulations in the outward facing conformation at equilibrium (last 50 ns) is also shown in **A**. The pairs are A79-T736 (ECL1-ECL3), G207-G850(TM3-TM9), G325-E968(ECL2-ECL4) at the extracellular gate, F71(TM1)-F728(TM7) pair and lower end Q343 (TM6)-Q986 (TM12) pair at the drug binding cavity, and K429-S1173 (site 1) and K1072-S528 (site 2) at the ATP-binding sites. In panel B and C, the protein initial structure (PDB: M41M, A) is shown as cartoon with half1 in pink and half2 in ice blue. The C_α_ of the landmark residues are drawn as green spheres.

On the other hand, the conformational changes at the NBDs were the most different between apo-Pgp and drug-occupied Pgp. In apo-Pgp, the conformational change achieved a closure at K429-S1173 (site 1) by −1.7 Å and an opening at K1072-S528 (site 2) by 4.1 Å with a final conformation of two equivalent sites (site1: 30.5 Å vs site2: 30.6 Å). On the other hand, in the presence of substrate, both site 1 (26.4 Å) and site 2 (24.2 Å) adopted a closure of −5.8 Å and −2.0 Å respectively. Both ATP binding sites K429-S1173 (site 1) and K1072-S528 (site 2) had significantly shorter distances in the drug-occupied Pgp than that of apo-Pgp. Since these pairs consist of residues from Walker A (K429/K1072) and LSGGQ (S1173/S528) motifs that directly sandwich ATP. The final pairwise distance was shorter in site 2 than site 1, which is also consistent with previous MD simulations 24.

### NBD distance and NBD alignment governed by lipids and drugs

The distances between the center of mass (COM) of each of the two NBDs were monitored over time (Fig. S5, Fig. S6). Apo-Pgp models (Fig. S5A) demonstrated larger oscillation and fluctuation than that of drug-occupied Pgp (Fig. S5B). However, all models eventually reached a stable COM distance by the end of the simulations (Fig. S5) and there was no significant difference (p=0.334) between apo-Pgp (39.5 Å) and drug-occupied Pgp (38.3 Å) (Fig. S6). Compared to the crystal structures, both apo-Pgp and drug-Pgp showed significant decreases of NBD COM distances by 5.4 Å (p=0.007) and 6.3 Å (p=0.006) but still over 10 Å away from a full NBD dimerization as in the outward facing conformation. However, the alignment angles and interfaces between the two NBDs were substantially different between apo-Pgp and drug-occupied Pgp simulations (Fig. S6, Fig. 3, Movie S1). Selected angles for measuring the alignment of NBDs were monitored using two sets of landmark residues on the ATP binding motif Walker A and LSGGQ: angle 1 was between K429-G1175-L1182 (site1) and angle 2 was between K1072-G530-L537 (site2). These angles are approximately 180° in canonical ATP binding for an NBD full-contact sandwich dimer outward-facing conformation (Fig. 3 B) as revealed by previous crystallographic data of an ABC transporter ^36^ and simulations ^33^. Therefore, the alignment of the two angles directly correlates to the favorability of high-affinity canonical ATP binding outward-facing conformation. The alignment angles in drug-occupied Pgp (Fig S6 B, C) are significantly larger than apo-Pgp at both site 1 (p=0.003) and site 2 (p=0.005). Compared to the crystal structure, apo-Pgp species showed significant decreased alignment angles at both site 1 (p =0.0099) and site 2 (p =0.0004) while no statistical differences between the drugs occupied Pgp and crystal structures at both site 1 (p=0.283) and site 2 (p=0.05). The simulations revealed that apo-Pgp models were significantly misaligned with an average of 112.9° at site 1 and 110.3° at site 2 while the drug-occupied Pgp demonstrate better alignment with average angles of 138.3° at site 1 and 140.6° at site 2.

**Fig 3.**
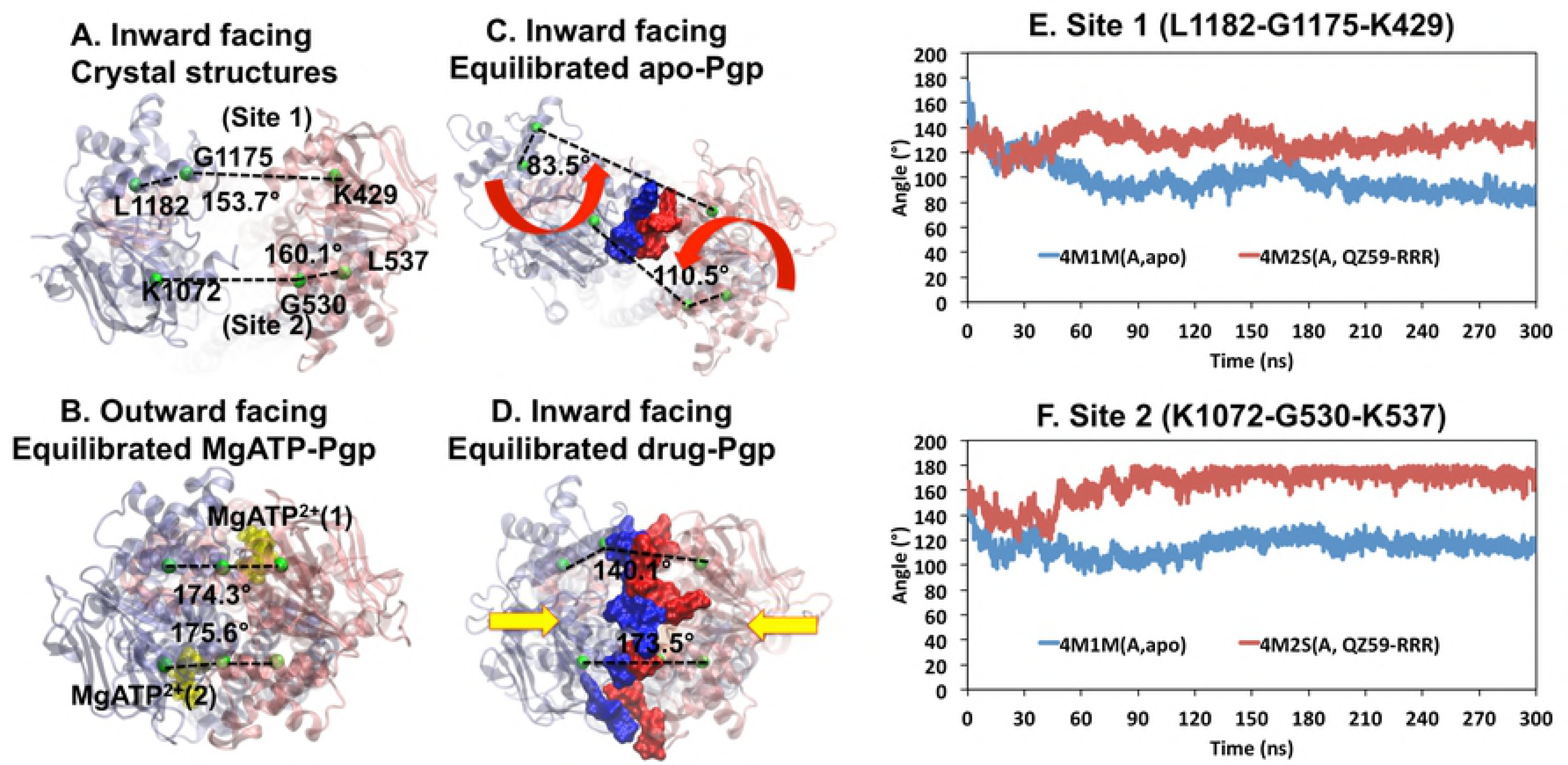
Differences between alignment angles and interfaces between apo-Pgp and drug-Pgp. The alignment angles (shown as black dashed lines) are monitored using the angles of C_α_ K429-G1175-L1182 (site1) and K1072-G530-L537 (site2). The proteins are shown as transparent cartoons with half1 in pink, half2 in ice blue. QZ59-RRR is shown as orange spheres and MgATP^2+^ as yellow spheres. The C_α_ of the landmark residues are drawn as green spheres. Red arrows show the directions of conformational changes. Interfaces (3 Å cut off) are drawn in surface with half1 in blue and half2 in red. **Panel A)** crystal structures 4M1M(mol A). **Panel B)** equilibrated structure of double MgATP2+ bound Pgp in the outward facing conformation at 200 ns 33. **Panel C)** equilibrated trajectory from 4M1M(A, apo) simulation at 300 ns. **Panel D)** equilibrated trajectory from 4M1M(A, QZ59-RRR) simulation at 300 ns.**E** shows the alignment angle at site 1 and **F** shows the alignment angle at site 2.

Since Pgp was crystallized *in surfo* in the absence of lipid bilayer, the effect of lipid bilayer interactions could not be gleaned from the structures. During our simulations reported here, lipid was observed to protrude through both sides (front and back) of the hydrophobic openings in Pgp called “bilayer portals”. Even though the linker region was not modeled in this simulation due to the unfolded nature in crystal structure, this region consisted of mostly polar and charged residues and highly hydrophilic made it very unlikely to interact with hydrophobic regions of TMDs. Instead, due to the hydrophobic chemical similarities between the middle regions of TMs and lipid molecules, such protrusion was also consistent with chemical principles. These interactions may explain differing NBD alignments between apo-Pgp and drug-occupied Pgp because the protrusions appeared to be different in both cases. The extent of portal opening was monitored using alignment angles between landmark residues (Fig. 4, S6, Movie S2). From the crystal structure *in surfo*, the front portal (between TM4 and TM6) has slightly smaller opening angle than the back portal (between TM10 and TM12) at 40.0° compared to 44.9°, respectively. However, in apo-Pgp simulations *in meso*, phospholipid exerted significantly different conformational effects (p=0.0137) on the two portals (Fig. 4A, B). The opening angle for the front portal increased to 48.0° and the back portal reduced to 36.8° (Fig. 4B). This conformational change is caused by lipid insertion between TM4 and TM6 (front portal) whereas lipids pushed TM12 closer to TM10 (back portal) rather than separating the two. The distinct difference between the front and back portal might due to the structural difference that TM12 has a break on the helix where as TM6 is completely helical (Fig. 5). However, drug-occupancy reversed this imbalance by similar portal angle (●p =0.474) on both sides (Fig. 4A, B) as shown for the front portal with an angle of 40.6° and the back portal at 40.3°, allowing an overall less lipid protrusion compared to apo-Pgp (Fig. 4A) and a more symmetric conformation similar to the crystal structure (Fig. 4A, B). The effect of lipids on the alignment angle was also analyzed visually for each case (Fig. S6). Lipid protrusion (Fig. S6) directly correlates to the NBD alignment angles (Fig. 3). For apo-Pgp (Fig. S5A, Fig. S6A-F), the more lipids protruded into the front portal, the greater the misalignment occurred. Fig. S6D, E shows minor protrusion that results in the best alignment angles in apo-Pgp (Fig. S6B).

**Fig 4.**
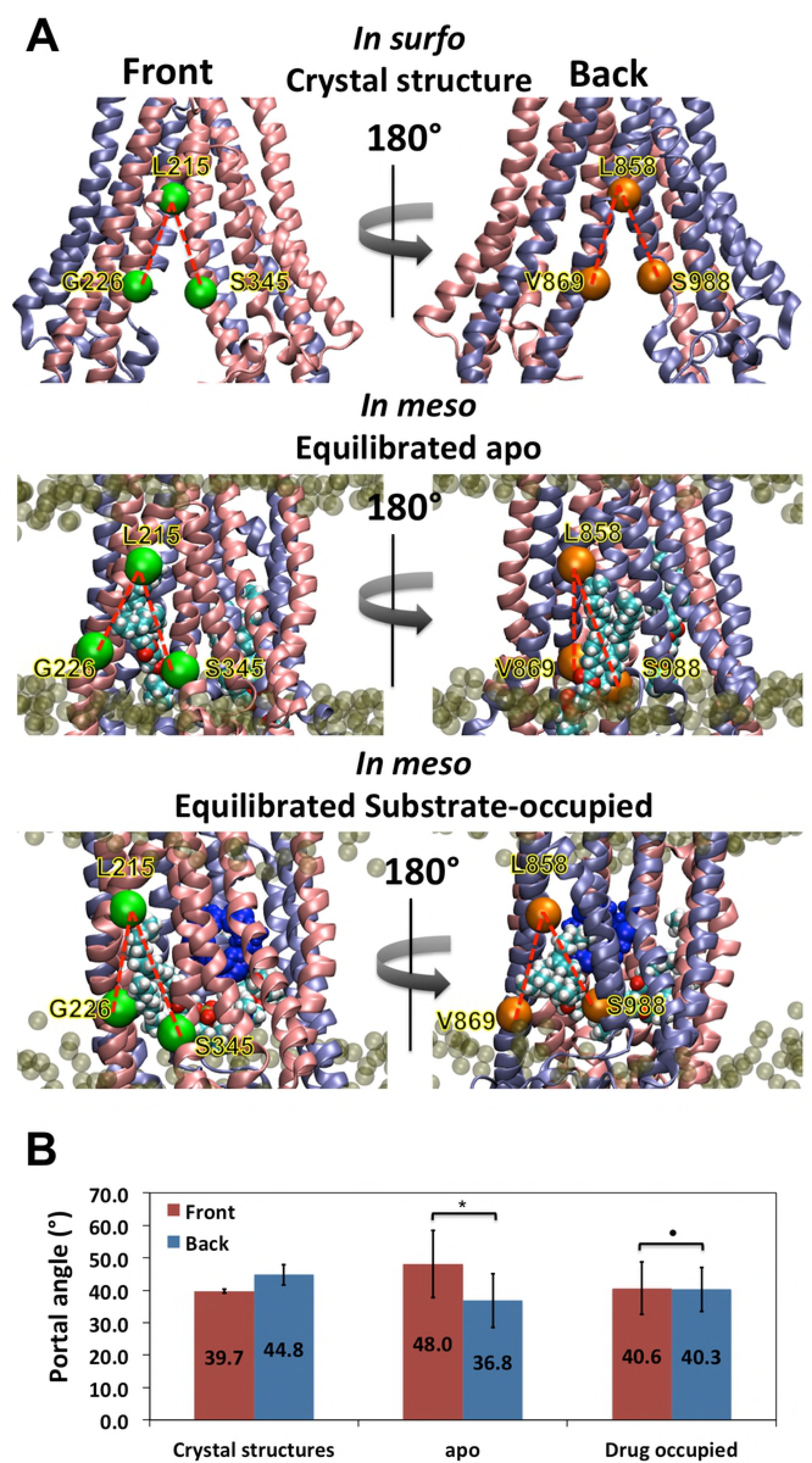
Influence of lipid and drug binding on conformation of Pgp bilayer portals. In Panel **A**, the bilayer portals opening angles (as red dash lines) are monitored using the angles of C_α_ G226-L215-S345 (Front) and V869-L858-S988 (Back). The proteins are shown as cartoon with half1 in pink and half2 in ice blue. QZ59-RRR drugs are drawn as blue spheres. The C_α_ of the front landmark residues are drawn as green spheres and the back landmark residues are drawn as orange spheres. Panel **B** shows the average angles over six crystal structures and averages of trajectories over the last 20 ns simulations for six apo-Pgp and six drug-occupied models. In apo Pgp, the difference between the front and back portals are significant (^*^p=0.0137) whereas in drug-occupied Pgp, no significant difference was observed between the front and back portals (●p=0.474).

**Fig 5.**
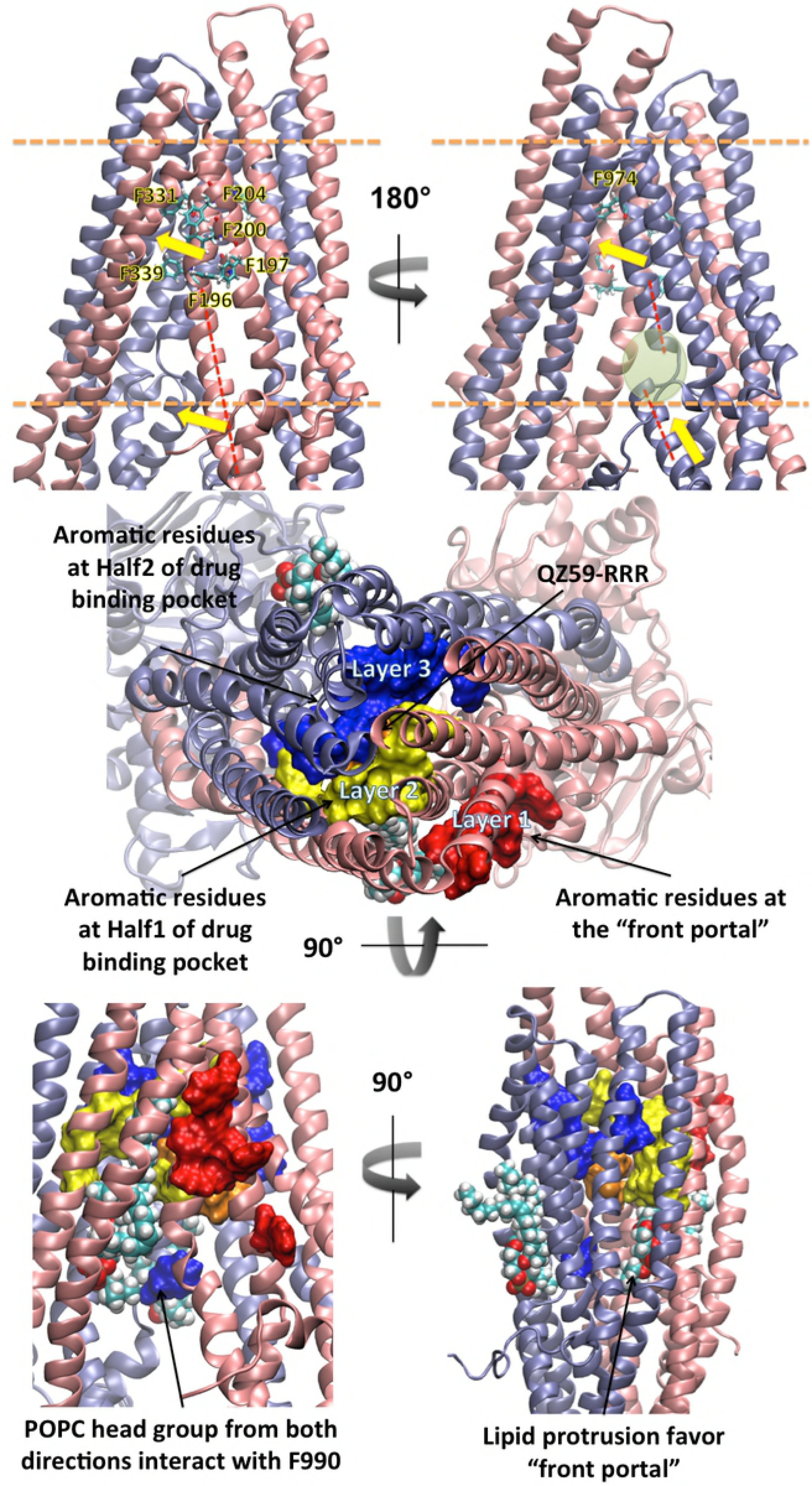
Bilayer portals exhibit distinct biochemical profiles. The “front portal” is formed by TMs 4 and 6, and the “back portal” is formed by TMs 10 and 12. In the top panel, the proteins are shown as cartoon with half1 in pink and half2 in ice blue. The phenylalanine residues are shown in licorice colored by elements. The boundaries of lipid bilayers are shown in orange dash lines. Helical axis of TM6 (Front) and TM12 (Back) are shown in red dash. Broken helix of TM12 is circled in green. Yellow arrows represent the directions of conformational change at TM6 and TM12. In the middle and bottom panels, aromatic residues are shown in surface with layer 1 in red, layer 2 in yellow and layer 3 in blue. Protruded lipids are shown in spheres colored by elements. Layer 1 aromatic residues in the front portal are directly exposed to the lipid bilayer, whereas Layer 3 aromatics near the back portal are located deeper into the internal drug-binding cavity.

The lipid intrusion separates TM4, 6 and TM10, 12. Since TM4/TM12 crosstalk with NBD2, and TM10/TM6 crosstalk with NBD1, such separations are transmitted from the TMDs to the NBDs and cause the misalignment of NBDs. On the other hand, drug occupancy provides a counter-force by interacting with TM6 and TM12, thus, counter-balancing the force of lipid protrusion and influencing the alignment angle for canonical ATP binding conformations. The central role of TM6 and TM12 in drug binding was also evidenced by previous biochemical experiments ^37^. Given that Pgp requires a lipid interface for substrate-stimulated ATPase activity and exhibits basal (drug-independent) ATPase activity, lipid protrusion and the observed NBD alignment must be reversible processes that would eventually allow alignment, ATP-binding and hydrolysis. Lipid induced misalignment of NBDs in the absence of drug occupancy may also function as a reversible kinetic brake to reduce needless ATP hydrolysis. The braking mechanism would conserve energy in a manner that insures more efficient consumption of ATP only when both lipid and drug are properly positioned in their respective interaction sites.

### Structural basis of drug recruitment preference through the front portal

POPC lipid head groups reached into the drug binding cavity in the absence of drug (Fig. S7 mol A, B, C, D and F) primarily from the “front portal”. In previous biochemical experiments ^38^, phosphatidylcholines (PCs), especially short chain PCs, were identified as transported substrates. Our observation of POPC intrusion into Pgp has two significant implications: 1) some lipids from the bilayer might gain full access to the internal cavity and be transported as a substrate, which may explain the so called “basal” ATPase activity of Pgp in the absence of drugs ^39^; and 2) since POPC intrusion occurred preferably through the front portal, other transported substrates might prefer accessing the internal cavity through the same portal.

It has been shown that hydrophobic substrates are absorbed by Pgp directly from the bilayer ^5^, likely via the portals that open to the inner leaflet. Significant biochemical differences between the two portals were apparent in our observations (Fig. 5). The front portal has 4 phenylalanine sidechains on TM3 (F196, F197, F200, F204) pointed toward the bilayer and 2 phenylalanine sidechains (F331, F339) on TM6 pointed towards TM5 and were also part of the drug-binding cavity. But, the portal in the back (half2) has only one phenylalanine (F974) on TM12 at the pseudo-symmetric position of TM6 (F331) and no other nearby aromatic sidechains pointing toward lipid. The counterpart residues at the back portal are L839, G840, I834, L847 and A941 none of which are aromatic but mostly with alkyl side chains similar to the lipid tails. TM6 and TM12 have very distinctive secondary structures in which TM6 is a continuous alpha-helix whereas TM12 has a break in the helix in the form of a coil loop in the inner leaflet region of the bilayer (Fig. 6, red dash lines and green circle). The gap on TM12 divides the helical “spring” into an upper half that connects to the drug binding cavity and the lower half that connects to NBD2. The broken helix should have different torque, elasticity, rotational range, and displacement vectors during Pgp conformational changes compared to that of the fully helical TM6. Indeed, TM6 and TM12 exhibited distinctly different (asymmetric) movements during simulations revealing a greater degree of flexibility for the upper and lower halves of the TM12 helical domains (Fig. 6, yellow arrow). As a result of such structural asymmetry, Phe 990 on TM12 at the lower boundary of the drug-binding cavity was observed accessible to lipid via the “front portal” whereas TM6 is not accessible to lipid near the “back portal”.

**Fig 6.**
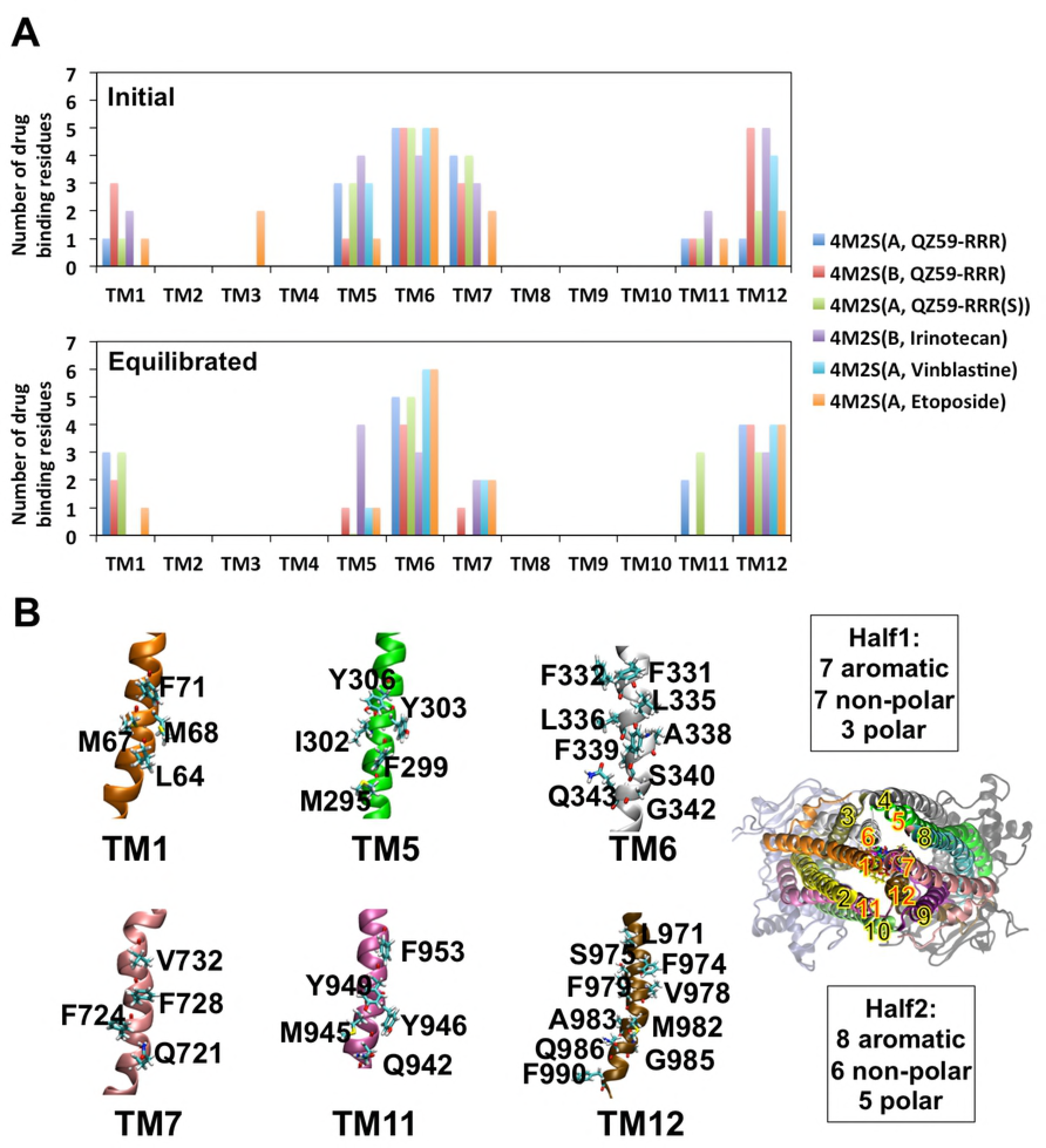
Poly-specificity of drug binding. Numbers of drug binding residues per TM are shown in **A.** Initial structures are crystal structure: 4M2S(A, QZ59-RRR), 4M2S(B, QZ59-RRR) and docking results: 4M2S(A, QZ59-RRR(S)), 4M2S(B, Irinotecan) and 4M2S(A, Vinblastine), 4M2S(A, Etoposide). Residues on six TMs that contribute to drug binding are shown in **B.** 3.0 Å cut off was used for ligand domain interactions with averaged over the equilibrated trajectories from the last 20 ns of 100 ns simulations. Drug molecules are shown in sticks in various colors.

The back portal is more aliphatic as with lipids, and the front portal has greater aromatic character. The Pi electron rich aromatic groups on phenylalanines would have distinct interaction chemistries with the drug including: 1) pi-pi stacking with pi electron rich regions, 2) cation-pi interaction with positively changed groups and 3) XH-pi (X=F, N, O) interaction with H-bond donors. A previous biochemical study^40^ involved a single mutation experiment on every phenylalanine on Pgp. The mutations of drug binding residues F331 (human F335) and F974 (human F978) were observed to reduce the Pgp drug resistance in a substrate dependent manner (F978A for all substrate where as F335A for half of the substrate), but all other portal aromatic residues (human F343, F200, F201, F204, F208) showed no significant effect on Pgp activity. The result suggested that due to the abundance of aromatic residues in the front portal region, the effect of a single mutation maybe rescued by compensatory functions provided by other aromatic residues.

Likewise, the Pgp drug-binding cavity itself has 15 aromatic residues (Fig. 6), which provides similar chemistries for attracting and binding Pgp substrates. The large aromatic components in the drug cavity thus assures the cavity to bind and translocate a wide range of aromatic and cationic compounds. Therefore, this enrichment of aromatic residues at the front portal indicates a potential preference for drug attraction to the drug than the back portal. In sum, there are three layers of aromatic residues (Fig. 5) around the drug binding cavity, layer 1 at the “front portal” where drug was first recruited, layer 2 and layer 3 contributed from half1 and half2 of Pgp where drug was sandwiched in between when recruitment is completed. Orientations of these aromatic residues and the pathway of lipid translocation as discussed above also indicates a dynamic relay of drug translocation: the drug interacts with layer 1 of aromatic residues on TM3, then shifts to layer 2 on TM6, eventually moving deeper into the drug binding cavity behind TM6 between layer 2 and layer 3. In contrast, the “back portal” lacks aromatic layers and lipid protrusion is much less observed, which further supports the suggestion that the “front portal” is the favored pathway for drug entrance. Therefore, layer 3 serves as “back wall” as opposite to “portal” because only the side that facing layer 2 showed drug/lipid interactions. The feature is reminiscence of the blocked back portal in the crystal structure of *C. elegans* PGP-1 ^41^ suggesting a conserved evolutionary bias against drug entry on this side of Pgp. We envision drug recruitment pathway as a single directional scheme (Layer 1 -> Layer 2 -> Layer 3).

### The structural basis of polyspecificity

Unlike conventional lock-and-key recognition between enzyme and substrate, Pgp accepts a large variety of drugs in a manner that is often termed as “polyspecific”. Pgp substrates, in general, often fail to obey Lipinski’s rules for efficacious drug potential in terms of having a larger size, greater hydrophobicity and greater number of rotatable bonds. Indeed, Pgp function may play a major role in defining the boundaries of efficacious drugs by reducing the bioavailability and increasing the clearance of these types of compounds. An alternative approach for studying substrate recognition and transport might then focus on synergistic domain mechanics rather than specific residues, bonds, etc. Conceptually, in order for a drug to be transported from one end of the binding cavity to the other, the drug cannot prefer solely one position during translocation. Instead, the drug may constantly move from one position to another to “respond” to the conformational change of Pgp. In other words, the allosteric regulation of drug binding may not occur through a particular residue or even a small number of residues, but rather, a synergistic effect from substrate interactions and changes between interactions with multiple TMs during protein conformational dynamics. MD simulations allow a dynamic inspection of the contribution of each TM to drug coordination(Fig. 6A). This study was based on the inward-facing conformation that drug molecules located in a relatively fixed location in the cavity due to the relatively fixed TMD conformation. Only, minor rotations and translations were observed for the drug molecules during simulation. From the comparison between the initial (crystal and docking structure) and last 20ns equilibrated trajectories, it is evident that only six TMs are involved in drug binding in the inward-facing conformation: TM1, TM5, TM6, TM7, TM11 and TM12. In particular, TM6 and TM12 are the primary binding domains with the most binding residues and were shared by all drugs throughout the entirety of all simulations. These observations are evidenced by both crystallography ^15^ and biochemical experiments ^40,42,43^. In one initial structure, Etoposide had a docking preference including some contact with TM3, but after 100ns simulation the drug equilibrated to a position where there was no contact with TM3. All other drugs demonstrated a dynamic binding network but maintained contact only with the six TM helices throughout the simulations (Fig. S8).

From the top view of Pgp (Fig. 7B), TM1, TM6, TM7 and TM12 form the core cavity for drug binding and TM5 and TM11 formed an extended cavity. The residues contributing to drug binding were analyzed as originally identified in the crystal structures (PDB 4M1M, 4M2S and 4M2T) and the equilibrated trajectories of six-drug binding models (Fig. 7B). Interestingly, the pseudo-symmetric domains do not have symmetric sequences in the cavity, resulting in a highly asymmetric binding cavity (Fig. 7B). Such asymmetry may also implicate the asymmetric allosteric regulation towards NBDs as discussed above. However, the total chemical components between the two halves are similar: “half1” has 7 aromatic residues, 7 non-polar residues and 3 polar residues whereas “half2” has 8 aromatic residues, 6 non-polar residues and 5 polar residues. In addition, residues of different types are evenly distributed on each domain, which provide a network of binding for a diverse population of drugs with different shape and size to bind. Such binding flexibility also assure the drugs be transported instead of retaining in the cavity. The simulations showed that drug binding was highly dynamic but that the number of binding residues on each TM is relatively stable during equilibrium.

**Fig 7.**
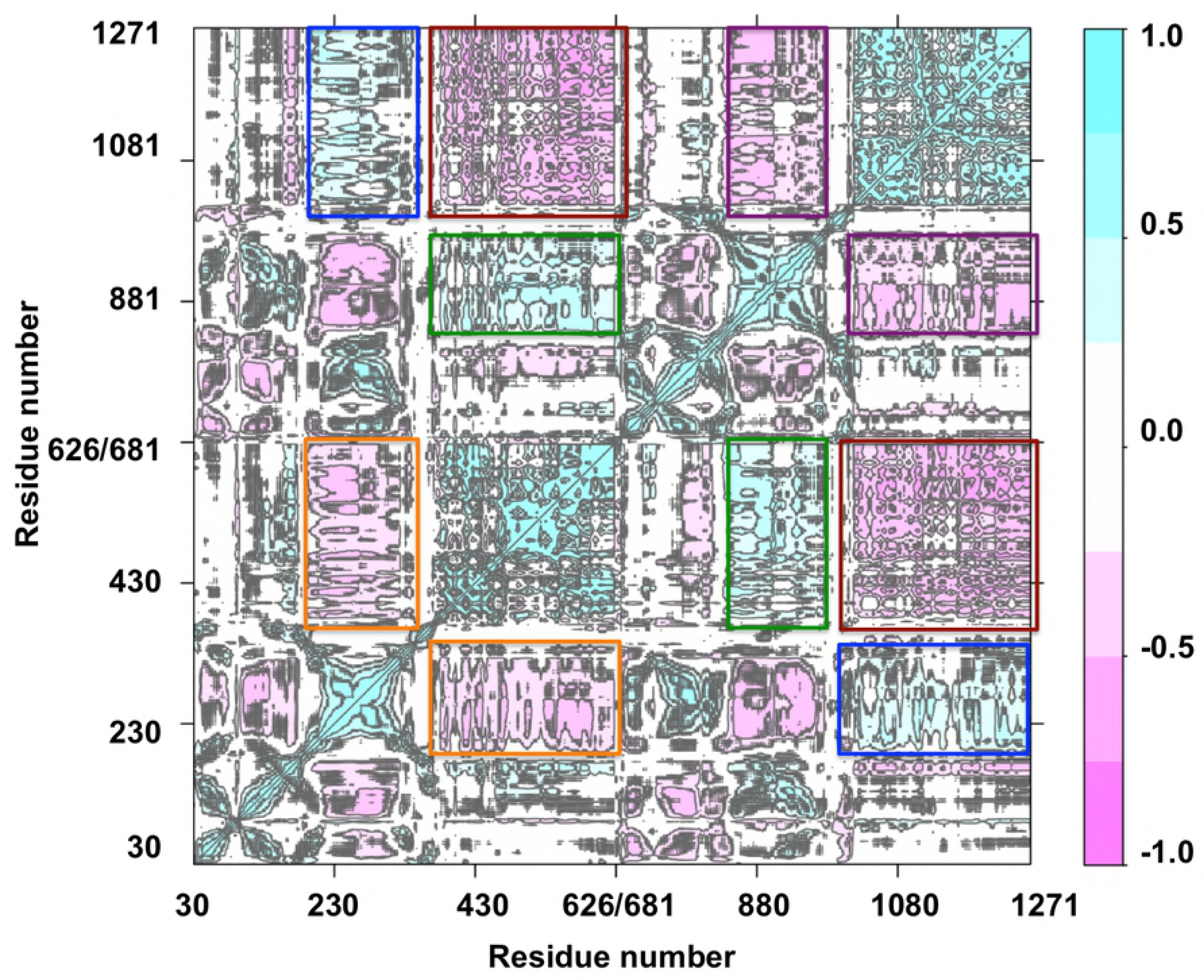
Dynamic cross correlation matrix (DCCM). DCCM was calculated using the alpha carbons in trajectories from 300 ns simulations of 4M2S(A, QZ59-RRR). DCCM of all simulations are shown in (Fig. S9). Squares highlighted the strong motion correlations between TMDs and NBDs. Red squares show strong negative correlations between NBD1 and NBD2. Orange squares show strong negative correlations between TM4-6 and NBD1. Purple squares show strong negative correlations between TM10-12 and NBD2. Blue squares show strong positive correlations between TM4-6 and NBD2. Green squares showed strong positive correlations between TM10-12 and NBD1.

### Allosteric regulation between NBDs and TMDs by DCCM analysis

In order to confirm the allosteric regulations between TMDs and NBDs discussed above, dynamic motion correlations between various residues and domains were analyzed with a dynamical cross-correlation matrix (DCCM) ^44^ on all Cα atoms (Fig. 7, S8). Positive correlations indicated movements along the same vector while negative correlations indicate movements in an opposite direction. Similar correlation patterns were observed in all simulations (Fig. S9). The result showed that NBD1 and NBD2 had strong negative correlation of motions (Fig. 7, red squares), which suggested that NBD1 and NBD2 moved in opposite directions and their motions are strongly synchronized. TM4-6 showed a strong positive correlation to NBD2 (Fig. 7, blue squares) while TM10-12 showed a strong correlation to NBD1 (Fig. 7, green squares) due to direct contact between these domains (TM4-5 with NBD2; TM10-11 with NBD1). In contrast, TM4-6 showed a strong negative correlation to NBD1 (Fig. 7, orange squares) and TM10-12 showed a strong negative correlation to NBD2 (Fig. 7, purple squares). Strong correlations between the motion of TMDs and NBDs were observed. As discussed above, TM4, 6 and TM10, 12 form portals where lipid protrusions were observed. In addition, TM6 and TM12 are major drug interacting domains. The analysis shows that the effects of lipids and drugs on TMDs could directly regulate the motion of NBDs.

### Normal mode analysis reveals the natural dynamics of Pgp

Lower frequency normal modes reflect the global natural movements of proteins whereas the higher frequency modes reflect regional movements. Fig. 8 depicts the motions of the first 5 lowest frequency modes (out of 3558 modes) of Pgp by normal mode analysis (NMA). Such analysis isolates and describes each motion separately whereas MD studies actually reveal the combinational effect of all thermal motions together. NMA also reveals how each domain cooperates with each other at one given frequency. Such a reduction improves understanding of conformational change in a more realistic way as opposed to an over simplified “open/close” description of Pgp. These motions were also observed during the conformational change of the simulations. For example, the misalignment of NBDs in the inward-facing conformation (Fig. 3C) is correlated to the movement of mode 1 and 2 in which two NBDs misalign by a counter-clockwise rotation (Fig. 8, Mode 1, 2) with lipid and without drug. NBD alignment in the outward-facing conformation (Fig. 3B) with lipid and without ATP is correlated to the movement of mode 5 in which the two NBDs exhibit a clockwise rotation (Fig. 8, Mode 5)^33^. Drug binding in combined with lipid promotes mode 3 and mode 4 with a more occluded site 2 compared to site 1. NMA also revealed asymmetric motions between two halves of Pgp, which indicates distinct structure-function relations between the two halves. The asymmetry of Pgp sequence thus determines the asymmetric nature of its structure and further dictates asymmetric dynamics.

**Fig 8.**
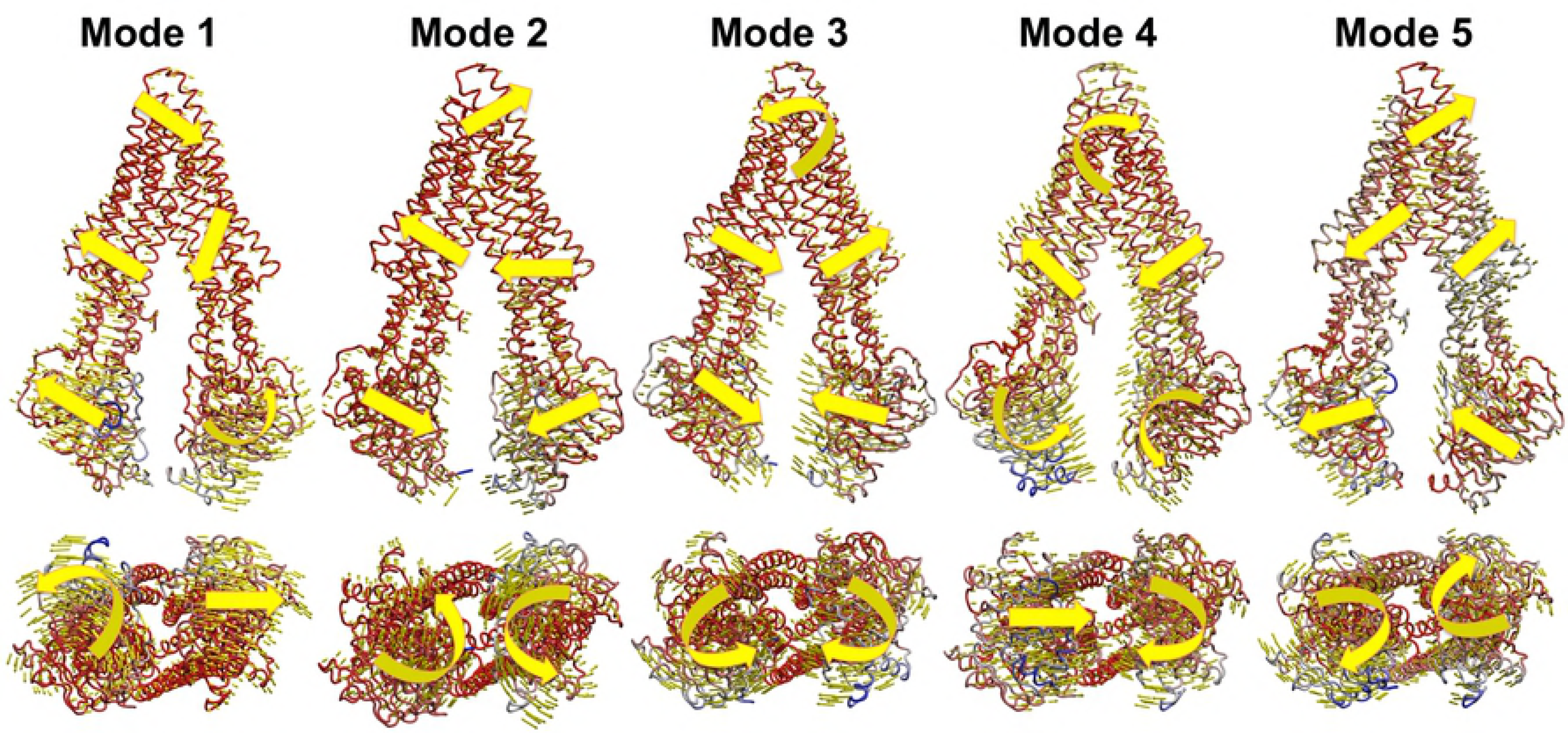
Normal mode analysis. Normal modes were calculated based on anisotropy network model algorithm ^45^ in ProDy program ^46^. The first 5 low frequency vibrational modes are shown in the Fig reveal the global thermal activity of Pgp. Protein structures are shown in color gradients with high frequency end in blue and low frequency end in red. Yellow arrows show the directions of thermal motions.

## Methods

### Docking and Modeling

Multiple apo- and drug-occupied mouse Pgp starting models obtained from the refined crystal structures were used in this study ^15^. Six apo-Pgp molecules were simulated using slightly different starting structures from PDB entries 4M1M (mol A and mol B), 4M2S (mol A and mol B) with ligands removed and 4M2T (mol A and mol B) with ligands removed. Two drug-occupied Pgp molecules from crystal structure 4M2S (mol A, mol B) with cyclic-tris-(R)-valineselenazole (QZ59-RRR) were used directly as initial structures for MD simulations. The crystal structures with left-handed ligands (4M2T) were not used because the ligands were not fully resolved in the crystal structures. In the structure of 4M2S (A), selenium (Se) atoms were substituted with sulfur (S) atoms as cyclic-tris-(R)-valinethiazole to simulate an additional model with a slightly different element component. In order to study the effects of other known drug substrates of Pgp, 30 different anti-cancer drugs and a Pgp modulator (Verapamil) were docked into the six apo-Pgp internal cavities using Autodock 4.2^47^ with a large grid covering the majority of the internal cavity (60×60×60 pixels) with a grid spacing of 0.375. Genetic algorithm parameters used for docking were as follows: number of evals=1,750,000; population size=150; runs=20 and rms tolerance for clustering=2.0 Å. Binding energies and clusters were analyzed (Table S1) and used to guide selection of compounds for MD simulation. The Pgp-drug complexes with the highest clustering score (Etoposide-4M2S-A) and the best binding energy score (Irinotecan-4M2S-B) and (Vinblastine-4M2S-A) were chosen. Etoposide-4M2S (A) had the highest clustering with 47 out of 100 hits in the same binding position (cluster is defined by rmsd ≤ 2.0 Å) and a good mean calculated binding energy of −10.3 kcal/mol. One conformation of Irinotecan-4M2SB and Vinblastine exhibited the best mean binding energy of −12.4 kcal/mol and −11.4 kcal/mol respectively. All the three drugs Irinotecan, Vinblastine and Etoposide have been shown to be *bona fide* substrates of Pgp in previous biochemical experiments ^48-50^. A total of twelve models with six apo-Pgp and six drug-Pgp models were thus simulated in this study as summarized in (Table 1). A 120 Å × 120 Å POPC lipid bilayer was modeled with Pgp protein inserted along the bilayer normal axis. In addition to matching the hydrophobic region of the TMDs to the hydrophobic region of the bilayer (Fig. S1A), biochemical studies revealed that Trp residues of membrane proteins near the lipid headgroup interface can form hydrogen bonds with the bilayer to “anchor” the protein with respect to the bilayer normal axis ^51^. This finding allowed us to place the -NH group on the indole ring of Trp (W208 and W851) in close proximity to the carbonyl groups on glyceride of the lipids (Fig. S1B, C), while the indole rings of other interfacial Trps (W44, W70, W132, W311, W694 and W704) were immersed in lipid without H-bonds. In order to achieve a proper lipid-protein interface for both leaflets of the lipid bilayer, Trp208, Trp311 and Trp851 were aligned with the glycerides of the outer leaflet of membrane whereas residues Trp44, Trp132, Trp228, Trp694 and Trp704 were aligned the glycerides of the inner leaflet of the membrane (Fig. S1A). After inserting protein into the bilayer using these criteria, overlapping lipids were removed with 0.6 Å cutoff distance from the protein. The membrane-embedded protein was then dissolved in a water box with a padding distance of >12 Å to every direction as standard periodic boundary. Physiological ion concentrations of 0.15 M NaCl and 0.002 M of MgCl_2_ were applied with system charge neutralized. The total number of atoms in the systems were ~209K.

### Force field

MD simulations utilized CHARMM22 with the CMAP correction of all-atom protein force field ^52^ for the protein section and CHARMM36 all-atom lipid force field ^53^ for the lipid section. Drug-like molecules including cyclic-tris-(R)-valinethiazole (QZ59-RRR(S), Se to S), and drugs such as Irinotecan, Vinblastine and Etoposide were parameterized using SwissParam that is suggested by the developers to be highly compatible with CHARMM force field ^54^. Due to lack of parameter for Se atom in the Swissparam data base (Merck Molecular ForceField and CHARMM General Forcefield), additional force field parameters for Se in cyclic-tris-(R/S)-valineselenazole (QZ59-RRR) were generated based on the density functional theory (DFT) calculations using B3LYP^55^ on 6-31+G^**^ basis set as used in CHARMM general force field^56^. Single point energy calculation and vibrational frequency calculation were carried after geometry optimization to acquire force constants of Se related parameters and partial charge information. The force field was tested to show normal structural and chemical behaviors of the drug molecules. Van der Waals parameters were imported directly from CHARMM general force field^56^ (Table S2). For both electrostatic and van der Waals calculations, 8 Å switching distance and a 12 Å cutoff distance were applied. Particle mesh Ewald (PME) method ^57^ was employed to calculate the long-range electrostatic interactions with 1.0 Å grid spacing. The TIP3P water model ^58^ was utilized in the simulations.

### MD simulations

All-atom MD simulations were performed for all models. The simulations were performed at isothermal–isobaric ensemble (NPT ensemble) at body temperature (310 K) and 1 atmosphere pressure. The constant temperature was maintained with Langevin dynamics using a damping coefficient of 1/ps. Constant pressure (1.01325 bar) was maintained using Nosé-Hoover Langevin piston pressure control ^59,60^ with the Langevin piston period of 200 fs and langevin Piston decay of 100 fs. A 2 fs time step was set for the entire simulation and trajectories were saved for every 5000 steps (10 ps). Multi-step pre-equilibrium simulations were performed before production phase. Each system was first subjected to conjugate gradient minimization for 10000 steps (20 ps). The system temperature was increased from 0 K to 310 K at a rate of 1 K/ps with the constraints (force constant k = 1 kcal/mol/Å^2^) on protein, ligands and the P atoms of lipids. An additional 5 ns simulation was performed at 310 K with the same constraints to equilibrate water. The constraints on P atoms of lipids were then removed for another 5 ns to equilibrate the lipid bilayer. Lastly, starting with an initial constraint (force constant k = 5 kcal/mol/Å^2^) on the protein backbone and ligands, the constraint was gradually removed at a speed of 0.5 kcal/mol/Å^2^ per 500 ps to accomplish equilibrated packing between the lipid and the protein. In the production phase, 100 ns to 300 ns simulations were performed without any constraint for the 12 models to observe potential conformation change and important meta-stable states.

### Computational Programs and data analyses

Docking experiments were carried out using Autodock4.2 ^47^. All atomistic MD simulations were performed using NAMD2.9 ^61^. DFT calculations were performed using Gaussian09 program ^38^. Normal mode calculation were based on anisotropy network model algorithm ^45^ in the ProDy program ^46^. Secondary structural analyses were based on the STRIDE algorithm ^62^. Dynamic cross correlation matrix (DCCM)^44^ was calculated using Bio3D^63^. Figures were created with PyMOL ^64^ and VMD ^65^. Statistical analyses were performed using Microsoft Excel. T-tests were performed for pairwise comparison. p <0.05 was regarded as significant difference.

## Disclosure and Acknowledgements

The authors declare no competing financial interests or conflicts of interest. We acknowledge funding support from NIH grant DP2 OD008591 and XSEDE award TG-MCB140067.

## Author Contributions

L.P. designed and performed the research, analyzed the data and wrote the paper. S.A. designed the research, analyzed the data and wrote the paper.

